# Epstein-Barr virus long non-coding RNA *RPMS1* full-length spliceome in transformed epithelial tissue

**DOI:** 10.1101/2021.02.07.430139

**Authors:** Isak Holmqvist, Alan Bäckerholm, Guojiang Xie, Yarong Tian, Kaisa Thorell, Ka-Wei Tang

## Abstract

Epstein-Barr virus is associated with two types of epithelial neoplasms, nasopharyngeal carcinoma and gastric adenocarcinoma. The viral long non-coding RNA *RPMS1* is the most abundantly expressed poly-adenylated viral RNA in these malignant tissues. The *RPMS1* gene is known to contain two cassette exons, exon Ia and Ib, and several alternative splicing variants have been described in low-throughput studies. To characterize the entire *RPMS1* spliceome we combined long-read sequencing data from the nasopharyngeal cell line C666-1 and a primary gastric adenocarcinoma, with complementary short-read sequencing datasets. We developed FLAME, a Python-based bioinformatics package that can generate complete high resolution characterization of RNA splicing at full-length. Using FLAME, we identified 32 novel exons in the *RPMS1* gene, primarily within the large constitutive exons III, V and VII. Two of the novel exons contained retention of the intron between exon III and exon IV, and a novel cassette exon was identified between VI and exon VII. All previously described transcript variants of *RPMS1* containing putative ORFs were identified at various levels. Similarly, native transcripts with the potential to form previously reported circular RNA elements were detected. Our work illuminates the multifaceted nature of viral transcriptional repertoires. FLAME provides a comprehensive overview of the relative abundance of alternative splice variants and allows for a wealth of previously unknown splicing events to be unveiled.

## Introduction

Alternative splicing brings considerable diversity to both human and viral transcriptomes. A number of elements including cassette exons, mutually exclusive exons, alternative splice sites and intron retention increases the variety of gene expression and regulation [1]. On the basis of short-read mRNA sequencing data from human tissues, it is estimated that around 95% of multiexon genes undergo differential splicing [2]. The utilization of RNA splicing in human herpesvirus transcription is highly dependent on the type of virus. With a few exceptions, the herpes simplex virus 1 genes do not contain introns and the mRNA maturation of the human genome is perturbed upon infection [3]. In contrast, multiple Epstein-Barr virus (EBV) genes contain variably included exons and are subjected to extensive splicing [4].

In EBV-associated neoplasms the virus expression is believed to be restricted to different types of latency programs. In EBV-associated gastric adenocarcinoma (GAC) and nasopharyngeal carcinoma, the most abundantly expressed poly-adenylated viral RNA is the 4 kilobases long non-coding RNA *RPMS1*, also referred to as BART (BamHI-A region rightward transcript) [5–8]. This high expression of *RPMS1* is also observed in the EBV-expressing cell line C666-1 [9]. *RPMS1* spans over 22 kilobases and contains seven main exons (I-VII) along with two minor cassette exons, Ia and Ib. Alternative exons within *RPMS1* as well as truncated versions of the transcript have been described [10, 11]. Particular segments of *RPMS1* can also backsplice and create circular RNA [12, 13]. Moreover, forty-four mature miRNAs (BART-miRNAs) are encoded within the introns of *RPMS1* and it has been proposed that the expression levels of these may be differentially regulated by alternating the splicing pattern [14, 15]. Furthermore, putative ORFs have been described from transcript variants of full-length and truncated versions *RPMS1* [16, 17]. However, the existence of these proteins have remained controversial.

Short-read RNA-sequencing provides the means to unbiasedly characterize transcriptional profiles with unprecedented speed, quantity and accuracy [18]. Nonetheless, comprehension of exon connectivity at single molecule level is irreversibly lost due to the fragmentation during the library preparation. In particular, this methodology fails to account for the relative abundance of alternatively spliced transcript at full-length resolution. Tools for high-throughput detection of alternative splicing in massive parallel sequencing data are readily available [19]. Splice-junction reads or paired-end mapping of coupled reads to two different exons allow for efficient detection of splicing, but these methods do not provide full-length resolution. In contrast, with the arrival of single-molecule sequencing technologies, it is now feasible to reveal the full spectrum of differential splicing [20]. However, the Oxford Nanopore Technologies long-read sequencing methodology is afflicted by the relatively low accuracy and high incidence of indels, which entails uncertainty in distinguishing novel splice sites from erroneous alignment [21, 22]. Bioinformatic tools such as FLAIR, which combine the strengths of the long-read and short-read sequencing data have proven to facilitate the correct alignment and annotation [23]. However, most tools are highly dependent on accurate reference annotations and may be limited in the allowance for overlapping exonic structures and would thus require laborious efforts for the detection of novel cassettes or alternative splice-patterns.

We have developed a bioinformatics tool package FLAME (Full-length Adjacency Matrix and Exon Enumeration) which in similar ways to previous tools combines long and short-read sequencing data. FLAME is a Python-based bioinformatics package that can generate complete high resolution characterization of RNA splicing at full-length single-molecule level. FLAME is especially designed to perspicuously account for the vast accumulation of previously undescribed alternative splicing events. By combining Nanopore long-read sequencing and publicly available short-read datasets of primary tumors, we set out to resolve the intricate splicing pattern of *RPMS1* in both an EBV cell line and a primary tumor by using FLAME.

## Results

The primary objective of the present work was to develop a bioinformatics pipeline capable of generating a compendious catalog of alternative splicing events. FLAME is a streamlined amalgamation of four main functions: (1) exonic filtering, (2) full-length variant enumeration, (3) adjacency matrix graphing and (4) novel splice site detection (Figure 1). These core functions are in turn constructed from several interconnected subfunctions (Table 1). Importantly, the novel splice site detection function allows for optional incorporation of corresponding short-read data to raise the dependability.

**Figure 1.**
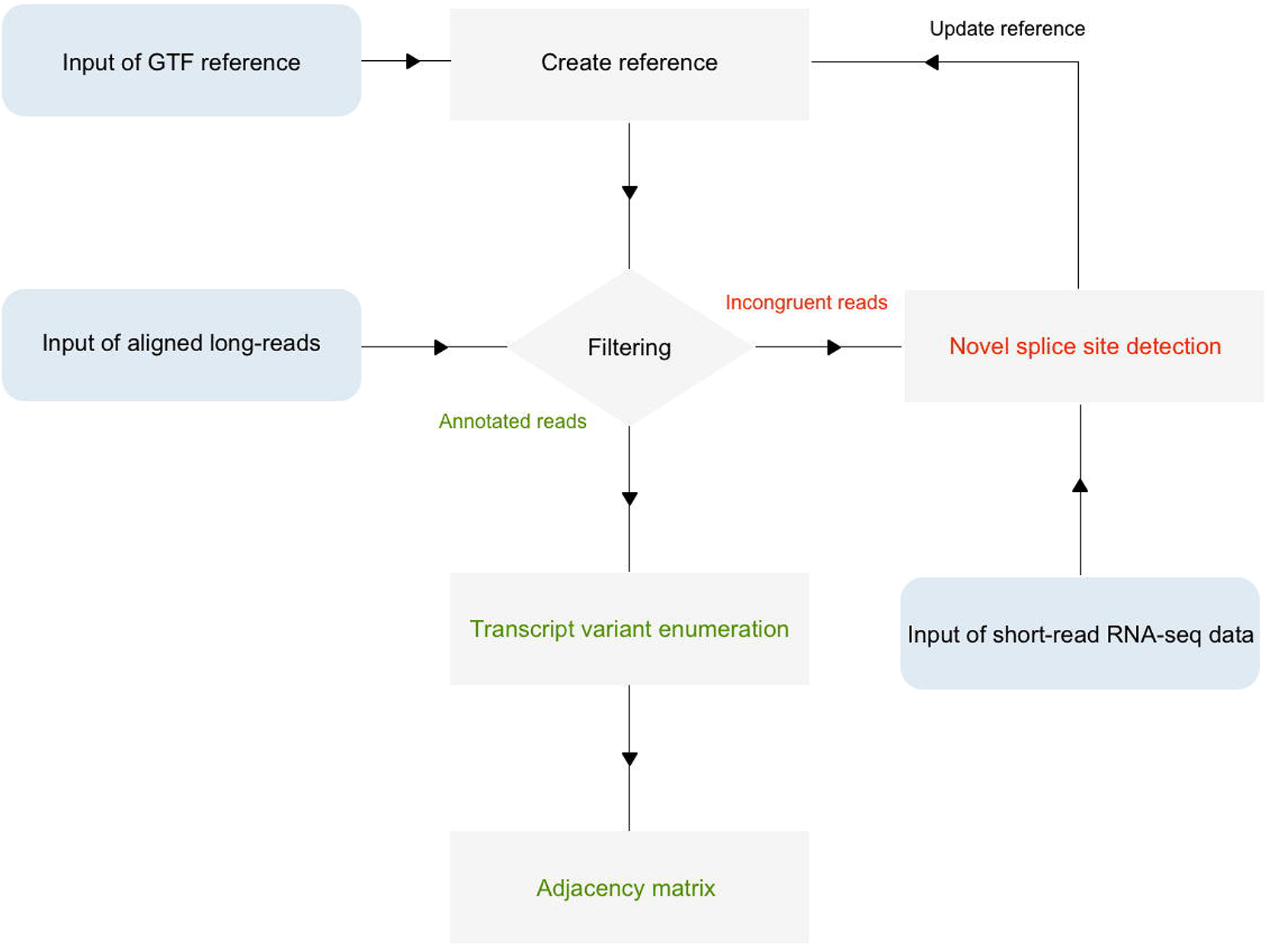
General organization of the FLAME pipeline. The ‘create reference’ function converts annotation written in GTF into a local variable for downstream use. Aligned long-reads are set against the reference in the subsequent ‘filter’ function. Here, transcript variants that contain incongruent exons are singled out and unannotated splice sites are catalogued. In the following ‘novel splice site detection’ function, unannotated splice site coordinates are quantified and set against a given FASTA format reference to scan for adjacent canonical dinucleotides (GU/AG). This step allows for optional incorporation of splice junction reads from corresponding short-read data to raise the fidelity. Novel exons that are deemed true upon manual inspection are then added to the reference variable for repeated cycling through the filter function. Transcript variants that are congruent with the reference annotation are passed on to an enumeration operation, in which the relative abundance is determined. The concluding adjacency matrix displays large-scale consecutive exon connectivity.

**Table 1:** Description of FLAME subfunctions.

| Core function | Subfunction | Description |
| --- | --- | --- |
| **Exonic filtering** | CREATEREF | Reads, filters and transforms the input reference file into a local variable. |
|  | FILTER | Categorizes reads as either *annotated* or *incongruent*. The filtering is performed on single-exon level and take into account the exon start position, end position and length. |
|  | TRANSLATEFUNC | Deciphers annotated exon ranges (in BED format) into a more straightforward nomenclature based on predefined exon designations from the local CREATEREF variable. |
| **Full-length isoform enumeration** | QUANTIFY | Collapses and quantifies transcript variants. |
| **Adjacency matrix graphing** | EMPTYADJMTX | Creates an empty adjacency matrix. |
|  | ANNOTATEDADJMTX | Creates a weighted adjacency matrix of consecutive exon connectivity. |
|  | INCONGRUENTADJMTX | Creates a weighted adjacency matrix of exon ranges retrieved from the FREQUENCYTHRESH function. |
| **Novel splice site detection** | INCONGRUENTSEPARATOR | Separates unannotated exon ranges into 5' and 3' splice sites. |
|  | FREQUENCYSITE | Extracts and transforms unannotated splice sites into frequency ranges in which the accumulative frequencies are plotted against gene positions. |
|  | FREQUENCYTHRESH | Extracts and returns unannotated splice sites based on a customizable threshold value. |
|  | SPLICESIGNAL | Locates unannotated splice sites and searches for adjacent dinucleotide splice site signals (GU/AG). |
|  | SHORTREAD | Scans the RNA-seq data to confirm the validity of unannotated splice sites. |

Total RNA was isolated from the nasopharyngeal carcinoma cell line C666-1 and a primary gastric adenocarcinoma (GAC) tumor tissue, in which EBV was detected by using RT-qPCR (Supplementary Figure 1). An adapted PCR-cDNA approach targeting *RPMS1* was subsequently employed to prepare full-length libraries for long-read sequencing. In total, 186,738 and 164,000 raw reads were generated from the C666-1 and GAC library respectively on an Oxford Nanopore Technologies MinION device and aligned to the EBV genome. Filtering based on the global start position of *RPMS1* eventually rendered 153,164 and 131,503 aligned reads, in the C666-1 and GAC dataset respectively. Inasmuch as PCR amplification and sequencing errors can overestimate alternative splicing events, we henceforth only considered transcript variants supported by ≥ 10 reads, referred to as intermediate-confidence. In addition, a high-confidence dataset with a threshold of ≥ 100 reads per transcript variant was implemented to more certainly rule out any technical artifacts. Given these threshold, only 11-13 % and 17-19 % was discarded in the intermediate-confidence and high-confidence datasets respectively, despite the high error rate of the long-read sequencing methodology (Supplementary Figure 2).

### The limitation of reliance on existing annotation

The initial `CREATEREF` function converts any genomic annotation in gene transfer format (GTF) into a local variable. The universal formulation of this function ensures that the program is applicable to both single- and multi-chromosomal conditions. As part of initializing `FLAME`, gene-specific annotation of *RPMS1* was accordingly extracted from NC_007605.1 by means of the `CREATEREF` function. However, this reference proved to be limited to describe the bulk of transcript variants as only 3.31% and 0.17% of the high-confidence reads from the C666-1 and GAC library respectively, was fully annotated. As expected, the novel splice site detection function immediately recognized exon II and the cassette exons Ia and Ib, which are all regarded as established amendments to the reference annotation. These three exons were therefore added to the reference annotation in order to achieve a reasonable baseline for further analysis. Utilizing this reference, the `FILTER` function retrieved 12.03% of the C666-1 reads, which were distributed on 22 intermediate-confidence transcript variants. As regards to the high-confidence dataset, four transcript variants were captured, representing 12.92% of the reads. When the GAC library was used as input, 17 intermediate-confidence transcript variants were captured, yet only representing 5.23% of the reads. Similarly, 6.24% of the reads, distributed on six transcript variants, were retrieved in the high-confidence dataset. In summary, this reference annotation failed to account for the overwhelming majority of *RPMS1* reads, which in and of itself constitutes a solid motive for recognition of alternative exons.

### FLAME singles out abundant unannotated exon boundaries from full-length transcripts

The novel splice site detection function was consequently initialized to sift through the large number of unannotated splice junction boundaries. FLAME was developed with the intention to provide a perspicuous means for de novo recognition of splice sites. The network of subfunctions implemented for this purpose revolves around three aspects: splice site usage frequency, canonical dinucleotides at the intron-exon junctions and incorporation of short-read reinforcement. These parameters are conveniently compiled in one output file. Publicly available short-read data from 106 EBV-positive nasopharyngeal carcinoma, 28 EBV-positive GAC tumors and one C666-1 dataset (equivalent to 459,854; 36,290 and 3,107 splice junction reads respectively) was accordingly incorporated via the SHORTREAD function (Supplementary Table 1). Thus, novel exonic elements had to meet the following criteria in order to be deemed authentic: 1) supported by ≥ 10 long-reads, 2) flanked by dinucleotide splice signals (GU)/AG and 3) both splice sites reinforced by complementary short-read data. By employing these criteria, which were all adequately systemized by means of the `INCONGRUENTSEPARATOR`, `INCONGRUENTADJMTX`, `FREQUENCYSITE`, `FREQUENCYTHRESH`, SPLICESIGNAL and `SHORTREAD` functions, `FLAME` distinguished 22 novel exonic elements in the C666-1 library and 30 exonic elements in the GAC library. The `FREQUENCYTHRESH` function was configured with a value of 1, meaning that only unannotated splice sites accounting for more than 1% of the incongruent reads were returned. In total, 32 novel elements were found, out of which 20 were common for both C666-1 and GAC (Figure 2). All exons were ultimately merged into one reference, which brought about an almost complete retrieval rate. Looking at the C666-1 intermediate-confidence dataset, 95.70% of the reads and 245 transcript variants were fully annotated, whereas 99.06% of the reads and 57 transcript variants were annotated in the high-confidence dataset. With respect to the GAC library, corresponding numbers in the intermediate-confidence dataset were 98,39% and 247 transcript variants, and 99,86% and 74 transcript variants in the high-confidence dataset (Supplementary Table 2).

**Figure 2.**
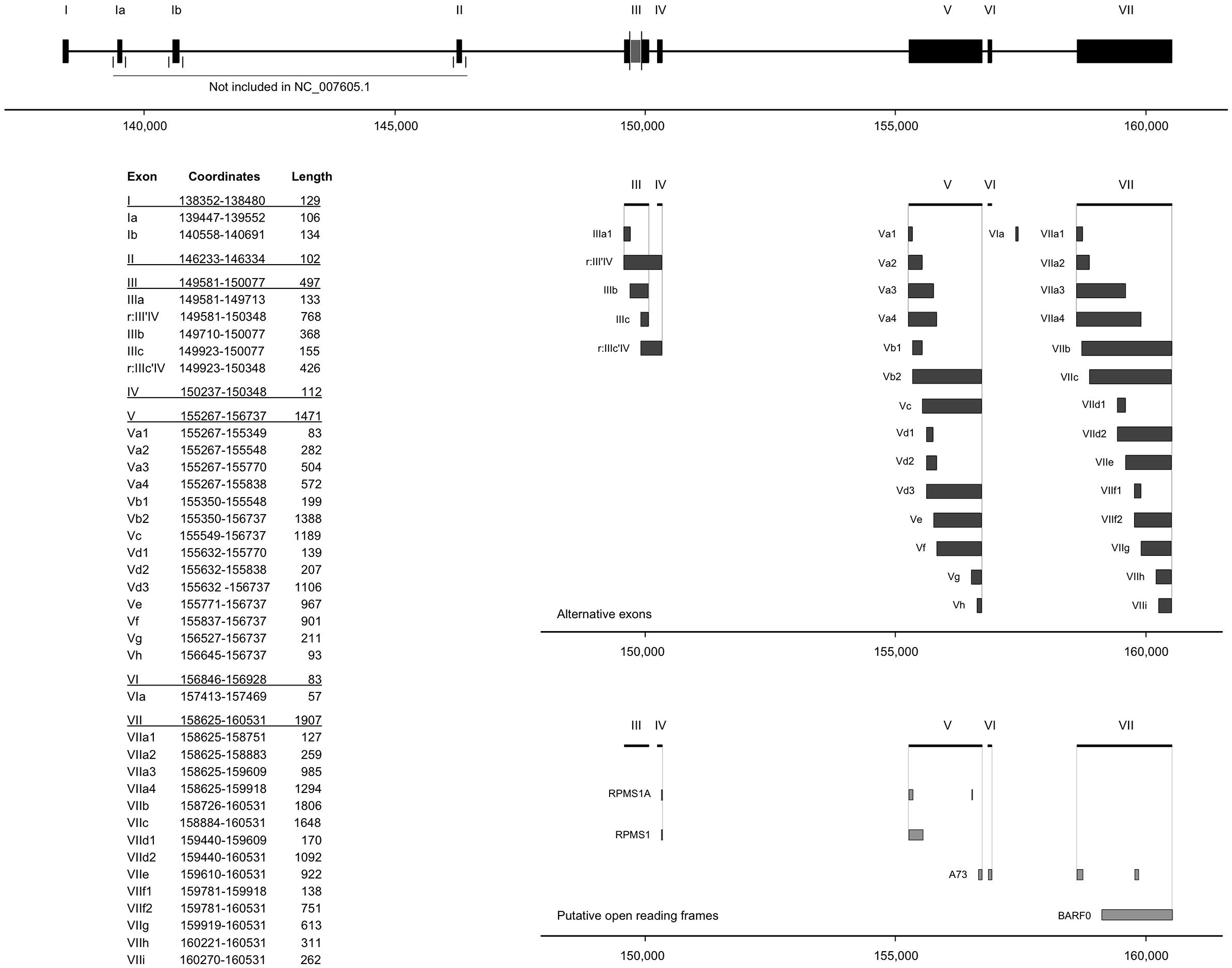
Comprehensive annotation of *RPMS1*. The upper segment displays the starting reference annotation of *RPMS1* as it is used in FLAME. Departures with respect to the RefSeq reference (addition of exon Ia, Ib and II) are indicated. Moreover, exon III is by convention regarded as one constitutive exon, although it is divided into two constituents in the RefSeq reference. Alternative exons discovered by FLAME are displayed in the middle segment. The nomenclature and coordinates of all exons are listed in the attached table. Putative open reading frames are displayed in the bottom segment. Numbers on the X-axis refer to the global position in the EBV genome.

### Benchmarking the performance

To assess the computational efficiency and robustness of the pipeline, we compared the performance of FLAME versus FLAIR using the GAC dataset with respect to retrieval of annotated reads, assembling of transcripts and runtime (Supplementary Table 3). Using FLAIR with the NC reference annotation, 549 reads (0.42%) were annotated and 11 full-length transcript variants were assembled on the basis of these. Despite expansion of the reference with all alternative exons obtained from the previous FLAME analysis, FLAIR was only able to annotate 4,602 reads (3.50%) and assemble 26 full-length transcripts compared with the 109,464 reads annotated as high-confidence by FLAME. However, 16 out of the 26 alternative transcripts assembled by FLAIR were lacking both exon V and VI, and six transcripts were missing exon III, IV, V and VI. This significant absence of otherwise established exons is notable. In addition, the intron between exon V and VI was retained in five cases. None of the transcript variants detected by FLAIR contained the canonical exon I-VII setup (Supplementary Figure 3). In comparison, using the same data, all of the transcript variants detected in FLAME, present in more than 1 % of the high-confidence dataset (see below), contained variants of the exon I-VII setup. The performance of FLAIR using the C666-1 dataset was comparable in all material respects, although the retrieval rate of annotated reads actually deteriorated when the reference annotation was expanded with alternative exons. The processing time for the analysis of *RPMS1* corresponded to 295 seconds and 54 seconds for FLAIR and FLAME respectively. Thus, FLAME performs alternative splicing analysis, including novel splice site detection, at 18.41% of the time it requires for FLAIR to finish its analysis.

### Characterization of alternative splice site usage in *RPMS1*

A novel cassette exon spanning over 57 bp and flanked with canonical dinucleotides was discovered within the intron between exon VI and exon VII. This exon, consequently designated exon VIa, was found to be mutually inclusive with exon VI and exon VIIa1 and occurred in three intermediate-confidence transcript variants in the GAC library. Notably, this exon was not detected in C666-1.

The `FILTER` function processes long-reads as aggregates of discrete exons; thus, reads are deemed as incongruent if merely one exon departs from the reference. However, a built-in variance function allows for a customizable window size of divergence with regard to exon global start position, global end position and length. It is worth to notice that this three-factor authentication entails awareness of overlapping structures and thereby allows for intron retention events and alternative splice sites within previously annotated exons to be categorized separately.

Exon III is divided into two segments in the reference annotation. By using FLAME, it was readily to draw the inference that exon IIIa and exon IIIc are virtually mutually inclusive. However, both the 5’ and 3’ splice sites of the alternatively spliced sequence in between are suboptimal since exon III was found to be fully retained in 64.30% of C666-1 high-confidence reads and 64.52% of high-confidence GAC reads. Moreover, an intron retention event spanning over exon III and exon IV (r:III’IV) was observed in two intermediate-confidence transcript variants (corresponding to 0.09% of the reads) in the GAC library. r:III’IV was however significantly more frequent in the C666-1 library as it appeared in seven intermediate-confidence transcript variants (1.57% of reads) and three high-confidence transcript variants (1.50% of reads). Furthermore, another intron retention event spanning from exon IIIc to exon IV was observed in one intermediate-confidence transcript variant (0.02% of reads) in the C666-1 library. The low percentage of the latter indicates that the pipeline gives the means for detection of extremely rare splicing events.

The emergence of alternative exons, which are defined here as novel splice sites in connection to previously known exons, was found to be restricted to longer exons, i.e. exon III, exon V and exon VII. Interestingly, however, all alternative splice sites appeared to be relatively weak in terms of inter-exonic connectivity as the outer boundaries of the exons were invariably engaged in splice junctions between constitutive exons. Figure 3 illustrates the relative strength of all acceptor and donor sites. For instance, the four discovered alternative donor sites within exon V are rarely consecutively connected to exon VI. Conversely, the seven alternative acceptor sites rarely connect with exon IV (Figure 3). When viewed collectively, the alternative splice sites within exon V were used in 33.12% of C666-1 high-confidence reads and 35.50% of GAC high-confidence reads. Notably, exon Vg comprise a major intra-exonic splice site acceptor in both GAC and C666-1 and provides an intrinsically strong donor site for consecutive coupling to the invariant exon VI. Eight alternative acceptor sites and four alternative donor sites were detected within exon VII. These were used in 27.36% of the high-confidence reads in C666-1. In stark contrast, the alternative splice sites were used in 68.18% of the high-confidence reads in the GAC library. Amongst the alternative exons within the boundaries of exon VII, VIIa1 in conjunction with VIIf2 was most abundant.

**Figure 3.**
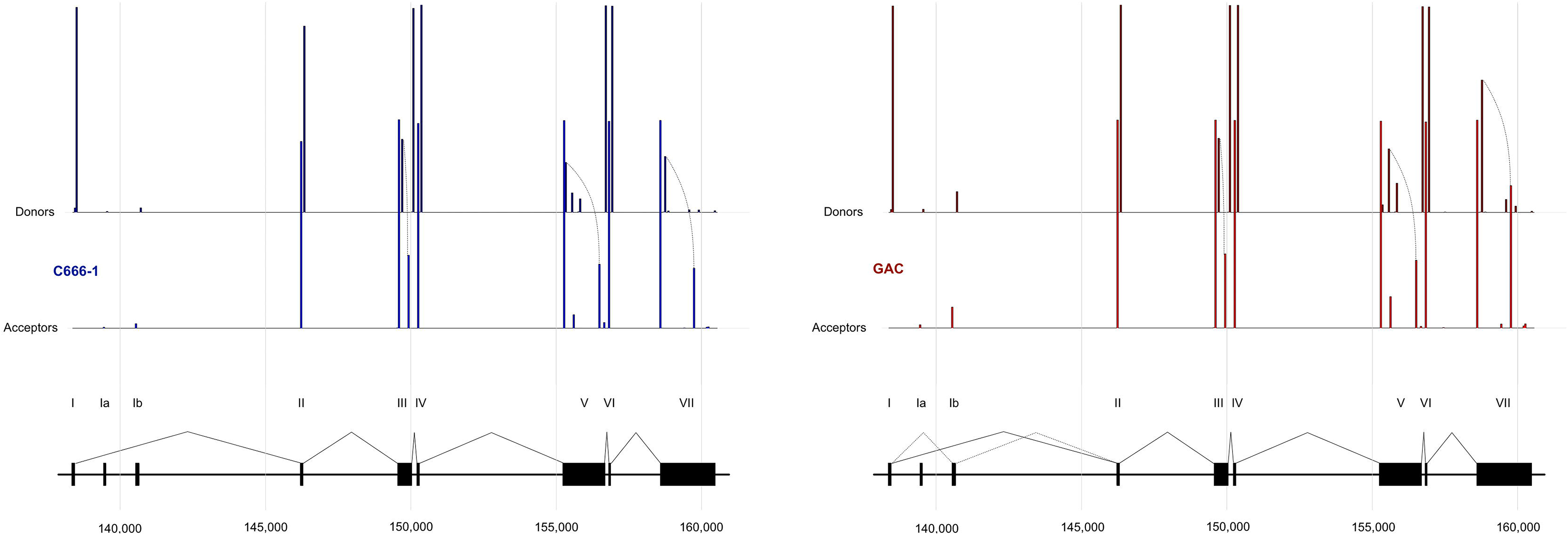
Splice site usage across *RPMS1*. Peaks corresponds to the usage frequency of a particular splice site. Dashed lines represent archetypical alternative splicing events within constitutive exons. The paramount pattern of inter-exonic splice junction connectivity is displayed in the bottom segment. Numbers on the X-axes refer to the global position in the EBV genome.

### Enumeration of alternative splicing at full-length resolution

The `TRANSLATEFUNC` function deciphers exon ranges into a more straightforward nomenclature based on predefined exon designations from the local reference variable. This is to avoid the laborious process of scrutinizing the sizable set of information in BED format. The subsequent `QUANTIFY` function basically collapses full-length permutations and hence allows for quantification of the relative abundance of transcript variants. The 15 most abundant splice forms of *RPMS1*, representing all variants supported by over 1.2% of high-confidence reads in the C666-1 library and 1% in the GAC library, are delineated in Figure 4. The 4.2 kb variant of *RPMS1* represents the paramount transcript in C666-1 and corresponds to 37.7% of high-confidence reads. It is worth to notice, however, that this splice form only represents 17.5% of the high-confidence reads in GAC (Figure 5). The palette of *RPMS1* transcript variants is largely attributable to alternative usage of multiple acceptor and donor splice sites within otherwise constitutive exons, i.e. exon III, exon V and exon VII. The repertory of splice forms is thereby shifted to shorter transcripts due to splitting of long exons, as illustrated in Figure 4. Differential usage of fully contained exons; that is cassette exons such as exon Ia and exon Ib, is of minor importance in this respect. A weighted adjacency matrix reveals overall patterns of consecutive exon connectivity in a quantitative manner (Figure 6). Vertical shifting within a column represents alternative acceptor sites for a given donor site, whereas horizontal shifting within a row represents alternative donor sites for a given acceptor site. In this way the full extent of alternative splicing within an exon, e.g. exon V, is comprehensively apprehended. Moreover, this graphical representation also conveys general indications regarding the exclusion rate of cassette exons, for instance exon Ia and exon Ib, exon II, and virtually mutually inclusive elements of split exons, e.g. exon IIIa and exon IIIc. Importantly, long-range exon connectivity possibly resulting from artifactual joining of primers appears in the top left corner.

**Figure 4.**
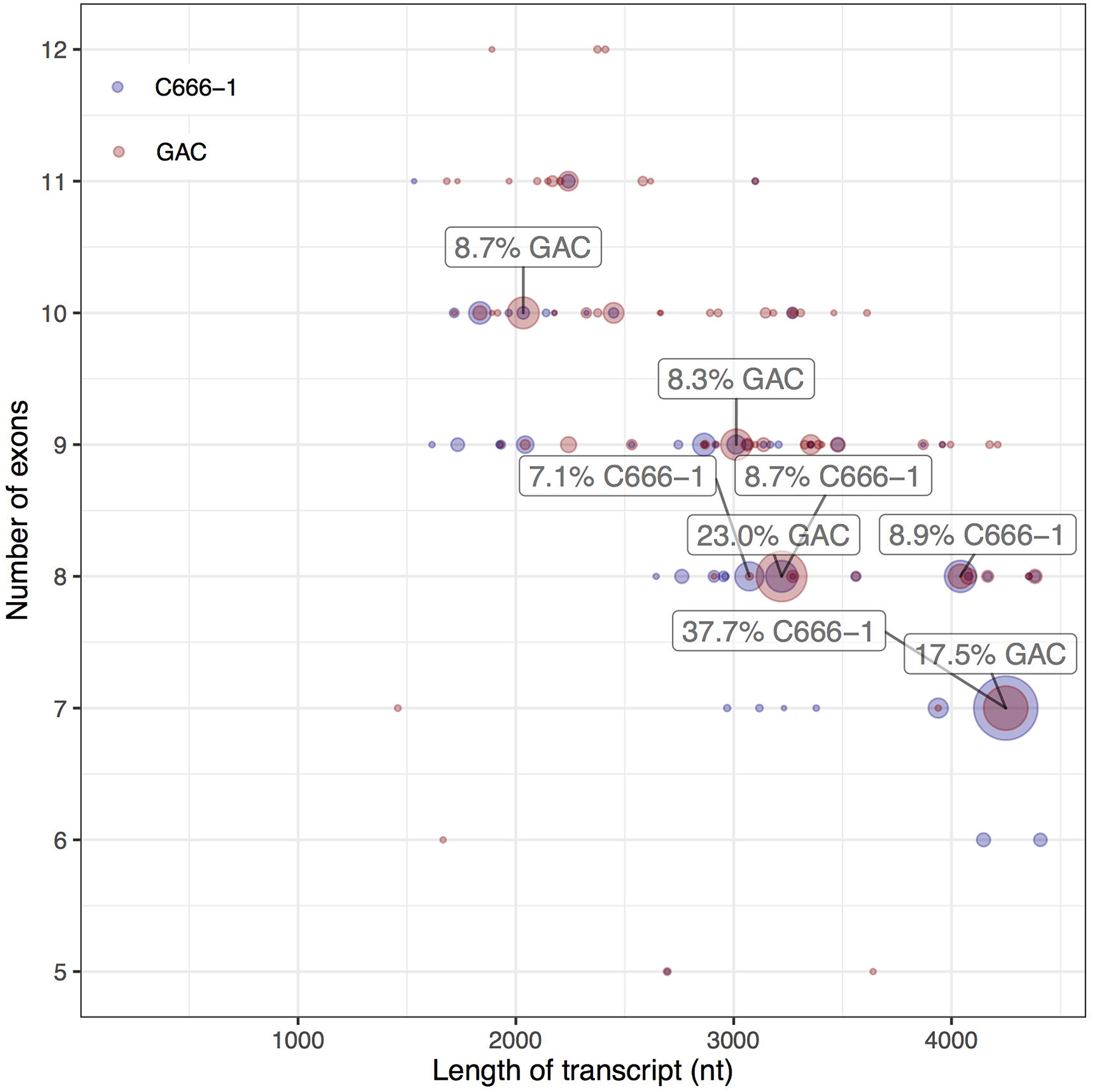
The most abundant transcript variants of *RPMS1*. The 15 most prevalent splice forms of *RPMS1*, representing all variants supported by ≥ 1.2% of high-confidence reads in the C666-1 library and 1% in the GAC library, are illustrated. Quantification of the relative abundance is expressed as the percentage of annotated high-confidence reads. Numbers on the X-axis refer to the global position in the EBV genome.

**Figure 5.**
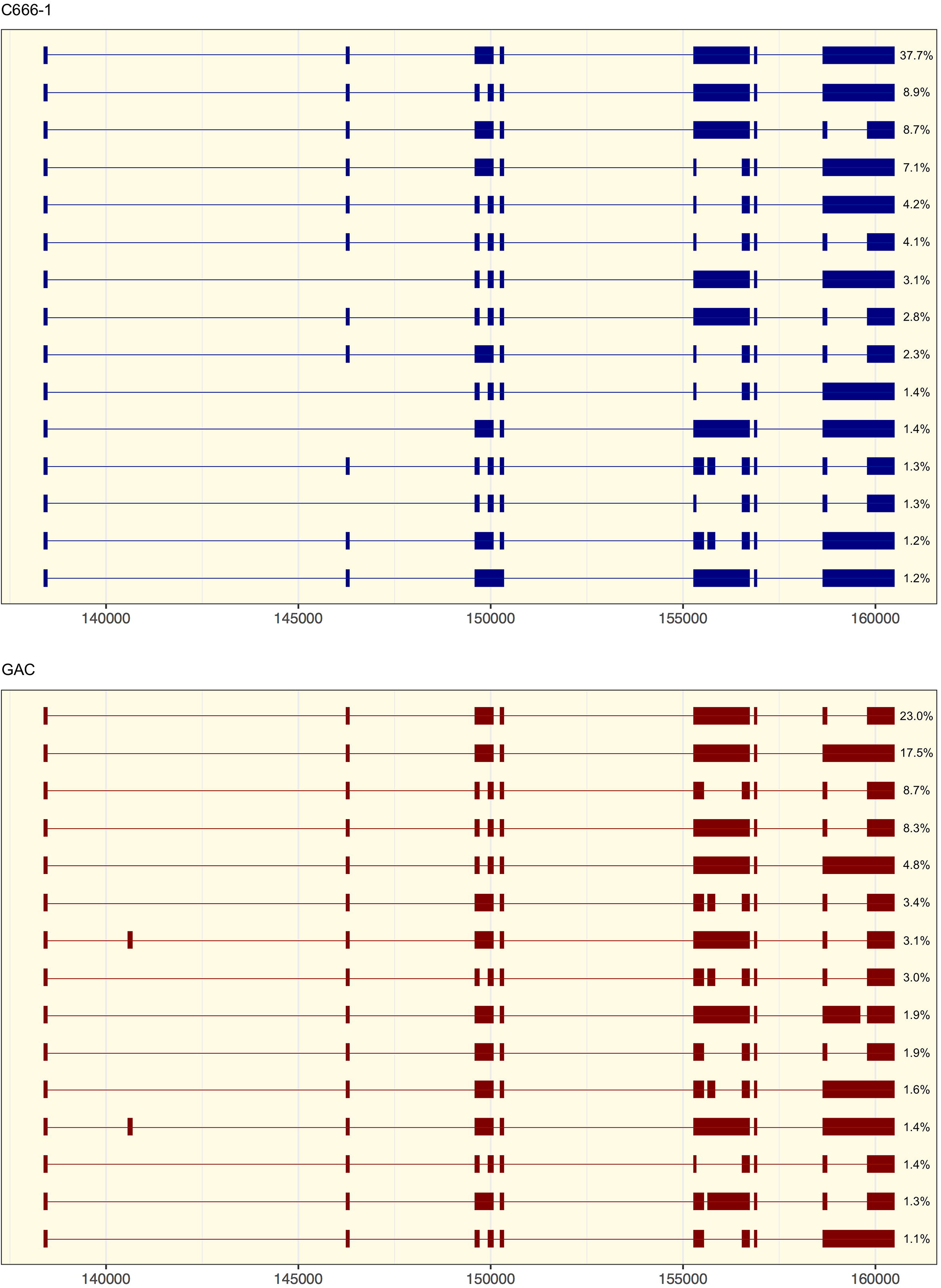
Diversity plot of high-confidence data. Each dot represents a unique transcript variant and the area correlates with the number of supporting reads expressed as percentage of fully annotated reads. The 4.2 kb splice form represents 37.7% and 17.5% of the reads in the C666-1 and GAC library respectively. Splicing within long exons pulls the center of density towards shorter transcripts with more exons. Transcript variants comprising < 5 exons (equivalent to 0.26% of the data) are not shown.

**Figure 6.**
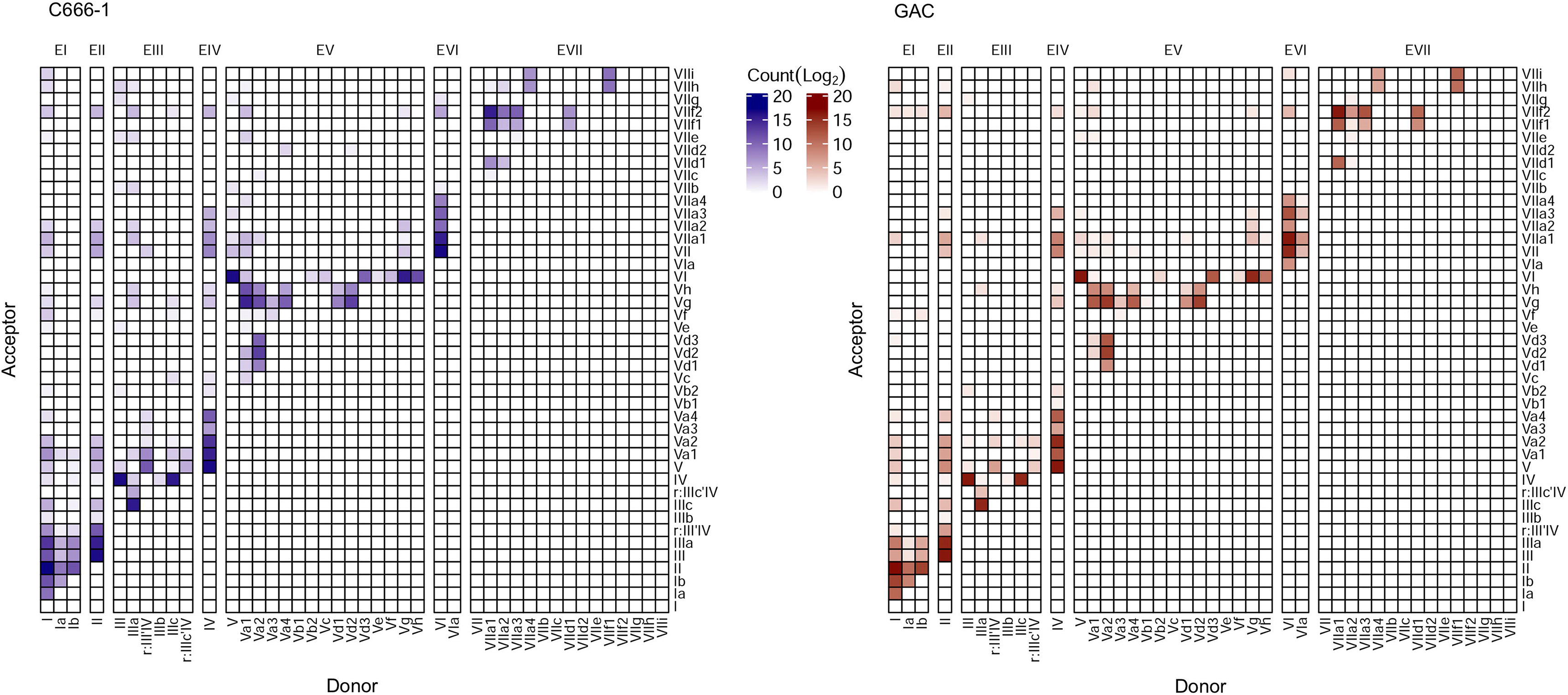
Weighted adjacency matrices outlining the overall patterns of consecutive exon connectivity in all long-read sequences. Vertical shifting within a column represents alternative acceptor sites for a given donor site, whereas horizontal shifting within a row represents alternative donor sites for a given acceptor site. Long-range exon connectivity possibly resulting from artifactual joining of primers appears in the top left corner.

### Functional implications of alternative splicing

The two most frequently observed circular RNA in primary tumors and cell lines derived from backsplicing from exon IV to exon II/III. The transcript variants with the potentiality to form circular RNA would therefore contain the splice donor of exon IV and the splice acceptor of exon II or exon III. The splice donor of exon IV was included in virtually all transcript variants in both datasets (99.70% of C666-1 and 99.59% of GAC high-confidence transcript variants). With the exception of extremely rare exon I to exon IIIc and exon II to exon IIIc splicing events, the splice acceptor of exon III was generally utilized. Likewise, the exon II was detected in practically all (99.55%) of the GAC high-confidence reads, but occasionally excluded in C666-1 transcripts. Moreover, nine circular RNA have previously been described to originate from exon VII. The splice donor for circularization in reactivated cells originates from the shared 3’ end of exon VIIa4 and exon VIIf1. In the GAC library, VIIa4 was found in three intermediate-confidence transcript variants, corresponding to 0.08% of reads; VIIf1 was on the other hand observed in 46 intermediate-confidence transcript variants and 9 high-confidence transcript variants, corresponding to 2.56% and 1.67% of reads, respectively. With regard to the C666-1 library, VIIa4 was found in 8 intermediate-confidence transcript variants, corresponding to 0.16% of reads, whereas VIIf1 was observed in 25 intermediate-confidence transcript variants and 3 high-confidence transcript variants, corresponding to 0.78% and 0.29% of reads, respectively. Two rare circular RNA with splice donor and acceptor within the intron between exon I and exon II previously detected in reactivated Akata cells were not detected in any transcript variants of GAC or C666-1. The absence and limited numbers of transcript variants with splice donors from the intron between exon I and exon II, and exon VIIa4/VIIf1 respectively imply that these transcripts mainly arise during reactivation or may be unique to B-lymphocytes, the conditions in which the circular RNAs were found.

The 15 most prevalent transcript variants constitute 82.49% and 86.64% of high-confidence annotated reads in GAC and C666-1, respectively. These transcripts contain 41 unique putative ORFs longer than five amino acids (Supplementary Table 4). Eight ORFs were only present in either of the libraries and the majority of these ORFs constituted less than five percent of the annotated reads. The suggested protein BARF0 (ORF_173) could be encoded from all transcripts in both GAC and C666-1. Previously described putative ORFs for RPMS1 (protein) and A73, but not RPMS1A (1.4%), were likewise found to be represented by a large fraction (60.2% and 54.1% respectively) of the annotated reads in GAC. In C666-1, from which most of these ORFs were discovered, RPMS1, RPMS1A and A73 were present in 63.8%, 20.4% and 20.5% respectively, of the annotated reads.

## Discussions

Long-read sequencing technologies have opened up the possibility to analyze full-length RNA transcripts splice variants in a high-throughput manner. The Oxford Nanopore Technologies is however limited by the relatively high error rates and current bioinformatic tools have not been able to categorize a significant proportion of the long-reads due to the complexity of RNA splicing. We have created a bioinformatic tool, FLAME, which combines short and long-read sequencing data to create a comprehensive and perspicuous description of any RNA transcript. The incorporation of complementary short-read data into the workflow reduces the deceiving impact of technical artifacts. Using FLAME we analyzed the complex spliceome of the EBV long non-coding RNA *RPMS1*. FLAME was compared with FLAIR and was shown to be five times faster and 24 times more efficient at annotating long-read sequences.

The spliceome of *RPMS1* shows that design of PCR primers for RNA without the detailed knowledge of the gene transcript variants may generate amplicons which neglects a large proportion of the transcripts. Furthermore, cell type/line and cell state may influence the spliceome. In the case of *RPMS1*, a state in which the EBV has been reactivated may lead to a significant perturbation of the spliceome, possibly due to the multiple protein encoding genes being transcribed in the opposite direction simultaneously. As previously shown, multiple *RPMS1* circular RNA were present only in a reactivated EBV B-lymphocyte cell line. Thereby the list of putative ORFs would also be highly influenced. With a comprehensive list of putative ORFs it would be possible to confirm or rule out the protein coding ability of *RPMS1* by unbiased approaches e.g. mass spectrometry. Currently, no known molecular function has been attributed to the long non-coding RNA *RPMS1*. Whether functional RNA domains exist within the *RPMS1* RNA remains to be seen, but our study has now added additional possibilities for creating secondary RNA structures.

FLAME has several aspects that are designed with the user experience in mind. It was written in a single programming language and designed to rely on as few external packages, software and tools as possible making FLAME very straightforward to use and almost self-sufficient in its implementation into different computational environments. Many facets of the program have been designed to allow for modularity, from the flexibility of implementing the user’s own threshold for the variance function and frequency analysis; to the ability to use each individual subfunction independently with either direct command-line interaction or through scripting. FLAME is applicable to both single-chromosomal organisms with complex gene expression like the *RPMS1* gene of the EBV and to multi-chromosomal organisms with complex gene expression. In this study we only considered transcript variants supported by at least ten or hundred reads to minimize the number of false positive variants created by technical artefacts or the erroneous sequencing technique while simultaneously report the most common transcripts. Depending on sequencing quality/depth and purpose of the study the frequency threshold can be adjusted to accommodate for the specific setting. Also, in order to detect microexons, an adjustment of the parameters needs to be made in order to narrow the window of long-read splice-junction detection. It should also be noted that a gene-specific PCR approach does not account for diversity pertaining to alternative promoter usage and/or alternative polyadenylation and thus needs to be complemented with an intricate knowledge of the specific gene in question. The present work provides a methodology for generating a compendious overview of the entire spliceome of any RNA transcript.

## Acknowledgement

We thank Dr. Ingemar Ernberg, Karolinska Institutet, Stockholm, Sweden, for the generous gift of the C666-1 cell line. We thank everyone involved in the Nicaraguan GAC sample collection initiative, especially Dr Lawrence Paszat, Dr. Reyna Victoria Palacios Gonzáles, and surgeons Dr. Alvaro Lopez and Dr. Carlos Ruiz. The results shown here are in part based upon data generated by the TCGA Research Network: https://www.cancer.gov/tcga. The computations were in part performed on resources provided by SNIC through Uppsala Multidisciplinary Center for Advanced Computational Science (UPPMAX) under project SNIC sens2018120. We thank the Bioinformatics Core Facility at the Sahlgrenska Academy for bioinformatics analyses. This study was supported by grants from Assar Gabrielssons Research Foundation, BioCARE National Strategic Research Program at University of Gothenburg, and the Wallenberg centre for molecular and translational medicine, University of Gothenburg, Sweden. The FLAME software is available on https://github.com/marabouboy/FLAME upon publication.

## Materials and Methods

### Cells

The nasopharyngeal carcinoma cell line C666-1 was grown in RPMI-1640 medium (Gibco) supplemented with 10% fetal calf serum and cultured at 37°C with 5% CO2. Total RNA was extracted using TRIzol reagent (Life Technologies) according to the supplier’s instructions. The eluate was subjected to DNase treatment (TURBO DNA-free™ Kit (Thermo Fisher Scientific) and then stored at −80°C.

### Patient samples

The gastric resection material was collected within a translational collaboration named “Immunological biomarkers for gastric cancer” (ethical approval number 2010/176-10). Samples were collected between June 2011 and July 2012 at Hospital Escuela Dr. Roberto Calderón Gutierrez (GHERCG), in Managua, Nicaragua [24]. During the study period, 15 patients were enrolled and biopsies from eleven patients were obtained. Punch biopsies were taken immediately after the resection and thereafter placed in RNA later and instantly snap-frozen. RNA was extracted with TissueLyser disruption using the RNeasy Mini Kit (QIAGEN). Five tumors were randomly selected and tested for EBV using RT-qPCR targeting *RPMS1* (Fwd: 5’-GATGTTTTGCGCCTGGAAGTTG; Rev: 5’-TCTCCTCGGACATCCAGTGTC) and *EBER-1* (Fwd: 5’-ACGCTGCCCTAGAGGTTTTG; Rev: 5’-AGACGGCAGAAAGCAGAGTC). *GAPDH* (Fwd: 5’-TCTCTGCTCCTCCTGTTCGA; Rev: 5’-GCCCAATACGACCAAATCC) served as an internal control.

### Long-read sequencing

First strand cDNA synthesis was primed with an oligonucleotide targeting the sequence immediately upstream of the poly-A signal (5’-TTGCATGTCTCACACCATGG). Approximately 2.5 μg of total RNA was incubated at 65°C for 5 min together with 20 pmol primer and 1 mM dNTP mix (Thermo Scientific), and thereafter instantaneously chilled on ice. The final reaction mixture was assembled in a total volume of 20 μl by adding 4 μl 5X RT Buffer (Thermo Scientific), 1 μl Maxima H Minus Reverse Transcriptase (Thermo Scientific) and 0.5 μl RNase OUT (Life Technologies), and incubated for 30 min at 55°C, 5 min at 85°C and then held at 4°C.

Full-length transcripts of *RPMS1* were selected by PCR amplification using Q5 High-Fidelity 2X Master Mix (New England Biolabs) according to the manufacturer’s protocol. A 2 μl-portion of the RT reaction mixture was carried into a total reaction volume of 25 μl and the following reaction was incubated at 98°C for 1 min prior to 18 cycles of [98°C for 10 s, 66°C 15 s, 72°C for 4 min], followed by a final extension at 72°C for 5 min and holding indefinitely at 4°C. Resulting amplicons were purified by incubation with 0.8X Agencourt AMPure XP beads (Beckman Coulter), followed by two washes with 200 μl of 75% ethanol and resuspension in 25 μl nuclease-free water. The remaining DNA concentration was measured with Qubit Fluorometer (Qubit DNA HS Assay Kit).

Subsequent end-prep with NEBNext Ultra II End repair/dA-tailing Module (E7546), Agencourt AMPure XP bead binding and Oxford Nanopore Technologies adapter ligation with NEB Blunt/TA Ligase Master Mix (M0367) was performed following the Direct cDNA Sequencing (SQK-DCS109) protocol version DCS_9090_v109_revJ_14Aug2019. The adapted and tethered library was enriched using 0.4X Agencourt AMPure XP beads washed with 2 x 200 μl Adapter bead binding buffer, and finally eluted in 14 μl Elution buffer (Oxford Nanopore Technologies).

The two libraries were separately loaded on FLO-MIN106D R9 flow cells according to the manufacturer’s specifications. The sequencing was performed on a MinION Mk1B device (MIN-101B) and operated through MinKNOW release 19.12.5. Raw data was basecalled using Guppy (3.6.1+249406c) configured with the high accuracy model (dna_r9.4.1_450bps_hac, default settings).

### Data processing

Two fast5 was generated from the Oxford Nanopore MinION, one from C666-1 and one from GAC, and was basecalled using the Guppy software from Oxford Nanopore Technologies (v3.6.1+249406c) using standard parameters to specify high-accuracy reads. Basecalling was throttled to only basecall 167,961 and 143,320 long-read sequences in fastq-format for the C666-1 and GAC samples respectively. This was done to have a comparable number of long-read sequences between the C666-1 and GAC sample. Long-read aware RNA aligner minimap2 (https://doi.org/10.1093/bioinformatics/bty191, v2.17-r941) was used to map the sequences with parameters of using the SPLICE preset of options and parameters while also specifying the exclusion of secondary alignment [25]. The NCBI RefSeq for EBV was used as the reference for the alignment for both samples. The generated SAM-files were sorted, indexed and compressed into the binary form using the samtools toolkit (https://doi.org/10.1093/bioinformatics/btp352, v1.10) [26]. The generated bam-files were filtered so as to remove reads categorized as supplementary and/or secondary reads. Further filtering on the bam-files were performed so as to require the inclusion of *RPMS1* exon 1. These filtering steps were done through an in-house bash-script. The filtered bam-files were then converted into bed12 format using the bedtools toolkit (https://doi.org/10.1093/bioinformatics/btq033, v2.26.0) [27]. These two bed12-files were then used as input for FLAME.

The C666-1, nasopharyngeal carcinoma and GAC bulk RNA-seq samples were retrieved from EMBL-ENA (ENA study accession number PRJNA501807 [28] and PRJNA397538 [29]) and TCGA (Samples that were classified as STAD and EBV positive [30]), respectively. The datasets were then preprocessed by Prinseq (https://doi.org/10.1093/bioinformatics/btr026, Version 0.20.3) and TrimGalore (https://github.com/FelixKrueger/TrimGalore, Version 0.4.4) [31]. The datasets were then aligned using STAR using the human HG38 (GRCh38) in fasta, the human annotation file in GTF format, the NCBI RefSeq EBV reference genome (NC_007605.1) in fasta format and the NCBI EBV annotation file in GTF format as reference. Specific parameters for all tools are available upon request.

FLAIR was used according to the developers’ instructions, using standard parameters for both the GAC and the C666-1 long-read RNA-seq, with each respective aforementioned shortread bulk RNA-seq pairing. FLAME was used with standard parameters for both the GAC and the C666-1 long-read RNA. The variance window was set at 20 nucleotides both upstreams and downstreams, and the novel splice site detection had a frequency threshold of more than 1 percent of the incongruent long-read sequences.

## References

1. Mollet, I.G., et al., Unconstrained mining of transcript data reveals increased alternative splicing complexity in the human transcriptome. Nucleic Acids Res, 2010. 38(14): p. 4740–54.

2. Pan, Q., et al., Deep surveying of alternative splicing complexity in the human transcriptome by high-throughput sequencing. Nat Genet, 2008. 40(12): p. 1413–5.

3. Tang, S., A. Patel, and P.R. Krause, Hidden regulation of herpes simplex virus 1 pre-mRNA splicing and polyadenylation by virally encoded immediate early gene ICP27. PLoS Pathog, 2019. 15(6): p. e1007884.

4. Farrell, P.J., Epstein-Barr Virus and Cancer. Annu Rev Pathol, 2019. 14: p. 29–53.

5. Raab-Traub, N., et al., Epstein-Barr virus transcription in nasopharyngeal carcinoma. J Virol, 1983. 48(3): p. 580–90.

6. Strong, M.J., et al., Differences in gastric carcinoma microenvironment stratify according to EBV infection intensity: implications for possible immune adjuvant therapy. PLoS Pathog, 2013. 9(5): p. e1003341.

7. Tang, K.W., et al., The landscape of viral expression and host gene fusion and adaptation in human cancer. Nat Commun, 2013. 4: p. 2513.

8. Li, A., et al., Transcriptional expression of RPMS1 in nasopharyngeal carcinoma and its oncogenic potential. Cell Cycle, 2005. 4(2): p. 304–9.

9. Verhoeven, R.J.A., et al., Epstein-Barr Virus BART Long Non-coding RNAs Function as Epigenetic Modulators in Nasopharyngeal Carcinoma. Front Oncol, 2019. 9: p. 1120.

10. Al-Mozaini, M., et al., Epstein-Barr virus BART gene expression. J Gen Virol, 2009. 90(Pt 2): p. 307–316.

11. Yamamoto, T. and K. Iwatsuki, Diversity of Epstein-Barr virus BamHI-A rightward transcripts and their expression patterns in lytic and latent infections. J Med Microbiol, 2012. 61(Pt 10): p. 1445–1453.

12. Toptan, T., et al., Circular DNA tumor viruses make circular RNAs. Proc Natl Acad Sci U S A, 2018. 115(37): p. E8737–E8745.

13. Ungerleider, N., et al., The Epstein Barr virus circRNAome. PLoS Pathog, 2018. 14(8): p. e1007206.

14. Marquitz, A.R., et al., Host Gene Expression Is Regulated by Two Types of Noncoding RNAs Transcribed from the Epstein-Barr Virus BamHI A Rightward Transcript Region. J Virol, 2015. 89(22): p. 11256–68.

15. Edwards, R.H., A.R. Marquitz, and N. Raab-Traub, Epstein-Barr virus BART microRNAs are produced from a large intron prior to splicing. J Virol, 2008. 82(18): p. 9094–106.

16. Smith, P.R., et al., Structure and coding content of CST (BART) family RNAs of Epstein-Barr virus. J Virol, 2000. 74(7): p. 3082–92.

17. Chen, H., et al., Expression of Epstein-Barr virus BamHI-A rightward transcripts in latently infected B cells from peripheral blood. Blood, 1999. 93(9): p. 3026–32.

18. Tang, K.W. and E. Larsson, Tumour virology in the era of high-throughput genomics. Philos Trans R Soc Lond B Biol Sci, 2017. 372(1732).

19. Jiang, W. and L. Chen, Alternative splicing: Human disease and quantitative analysis from high-throughput sequencing. Comput Struct Biotechnol J, 2021. 19: p. 183–195.

20. Garalde, D.R., et al., Highly parallel direct RNA sequencing on an array of nanopores. Nat Methods, 2018. 15(3): p. 201–206.

21. Dohm, J.C., et al., Benchmarking of long-read correction methods. NAR Genomics and Bioinformatics, 2020. 2(2).

22. Byrne, A., et al., Nanopore long-read RNAseq reveals widespread transcriptional variation among the surface receptors of individual B cells. Nat Commun, 2017. 8: p. 16027.

23. Tang, A.D., et al., Full-length transcript characterization of SF3B1 mutation in chronic lymphocytic leukemia reveals downregulation of retained introns. Nat Commun, 2020. 11(1): p. 1438.

24. Thorell, K., et al., In Vivo Analysis of the Viable Microbiota and Helicobacter pylori Transcriptome in Gastric Infection and Early Stages of Carcinogenesis. Infect Immun, 2017. 85(10).

25. Li, H., Minimap2: pairwise alignment for nucleotide sequences. Bioinformatics, 2018. 34(18): p. 3094–3100.

26. Li, H., et al., The Sequence Alignment/Map format and SAMtools. Bioinformatics, 2009. 25(16): p. 2078–9.

27. Quinlan, A.R. and I.M. Hall, BEDTools: a flexible suite of utilities for comparing genomic features. Bioinformatics, 2010. 26(6): p. 841–2.

28. Edwards, R.H., R. Dekroon, and N. Raab-Traub, Alterations in cellular expression in EBV infected epithelial cell lines and tumors. PLoS Pathog, 2019. 15(10): p. e1008071.

29. Zhang, L., et al., Genomic Analysis of Nasopharyngeal Carcinoma Reveals TME-Based Subtypes. Mol Cancer Res, 2017. 15(12): p. 1722–1732.

30. Cancer Genome Atlas Research, N., Comprehensive molecular characterization of gastric adenocarcinoma. Nature, 2014. 513(7517): p. 202–9.

31. Schmieder, R. and R. Edwards, Quality control and preprocessing of metagenomic datasets. Bioinformatics, 2011. 27(6): p. 863–4.

